## Supplementary material for "The gut commensal *Blautia* maintains colonic mucus function under low fiber consumption through short-chain fatty acid-mediated activation of Ffar2": S1

### Supplementary Figure 1

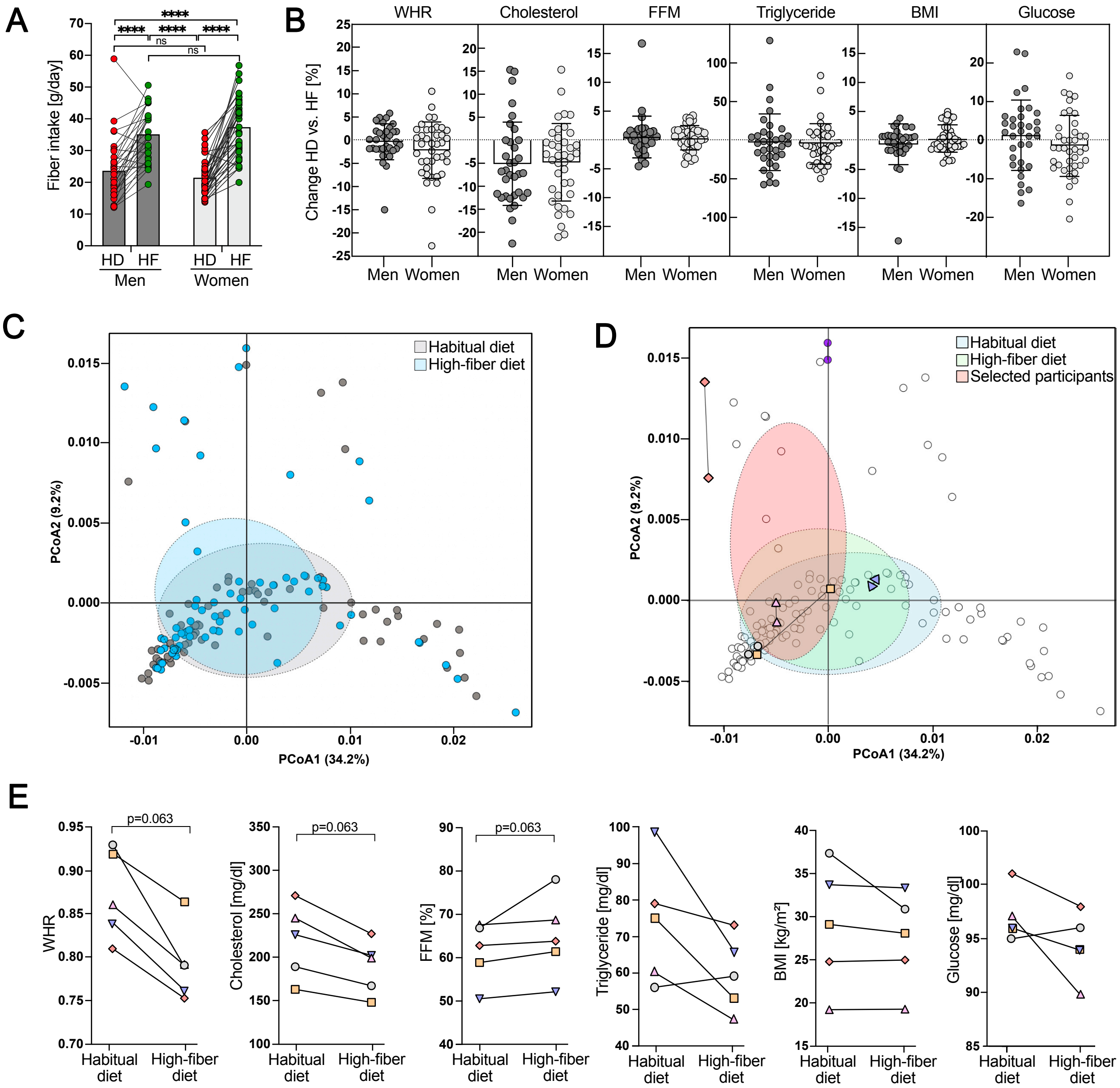


*Supplementary Figure 1*: (A) Fiber intake of male and female participants during habitual diet (HD) and after 12 weeks of high fiber (HF) intervention. (B) Change in metabolomic parameters for men and women after HF intervention. (C) Stool bacteria composition (Weighted UniFrac) of the study participants on their HD and after the HF intervention. (D) Participants selected for microbiota transplantations are highlighted in coloured symbols and indicated with a red cloud. Please note that ◆ was included only in the first FMT experiment (Fig.1) while ● was only included in the second FMT experiment (Fig 3). (E) Metabolic parameters of the 5 individuals selected as donors for FMT. Normal distribution of the data was tested using D'Agostino-Pearson test (A, B, E). Statistical significance was tested using 2-way ANOVA with Tukey's multiple comparison test (A), an unpaired t-test for normally distributed data and Mann-Whitney test for non-normally distributed data (B), PERMANOVA with 999 permutations (C, D) and Wilcoxon matched-pairs signed rank test (E) with p<0.0001(****) considered statistically significant. WHR=waist-to-hip ratio, FFR= free fat mass, BMI= body-mass index. Linked to Figure 1.
