## Supplementary material for "The gut commensal *Blautia* maintains colonic mucus function under low fiber consumption through short-chain fatty acid-mediated activation of Ffar2": S2

### Supplementary Figure 2

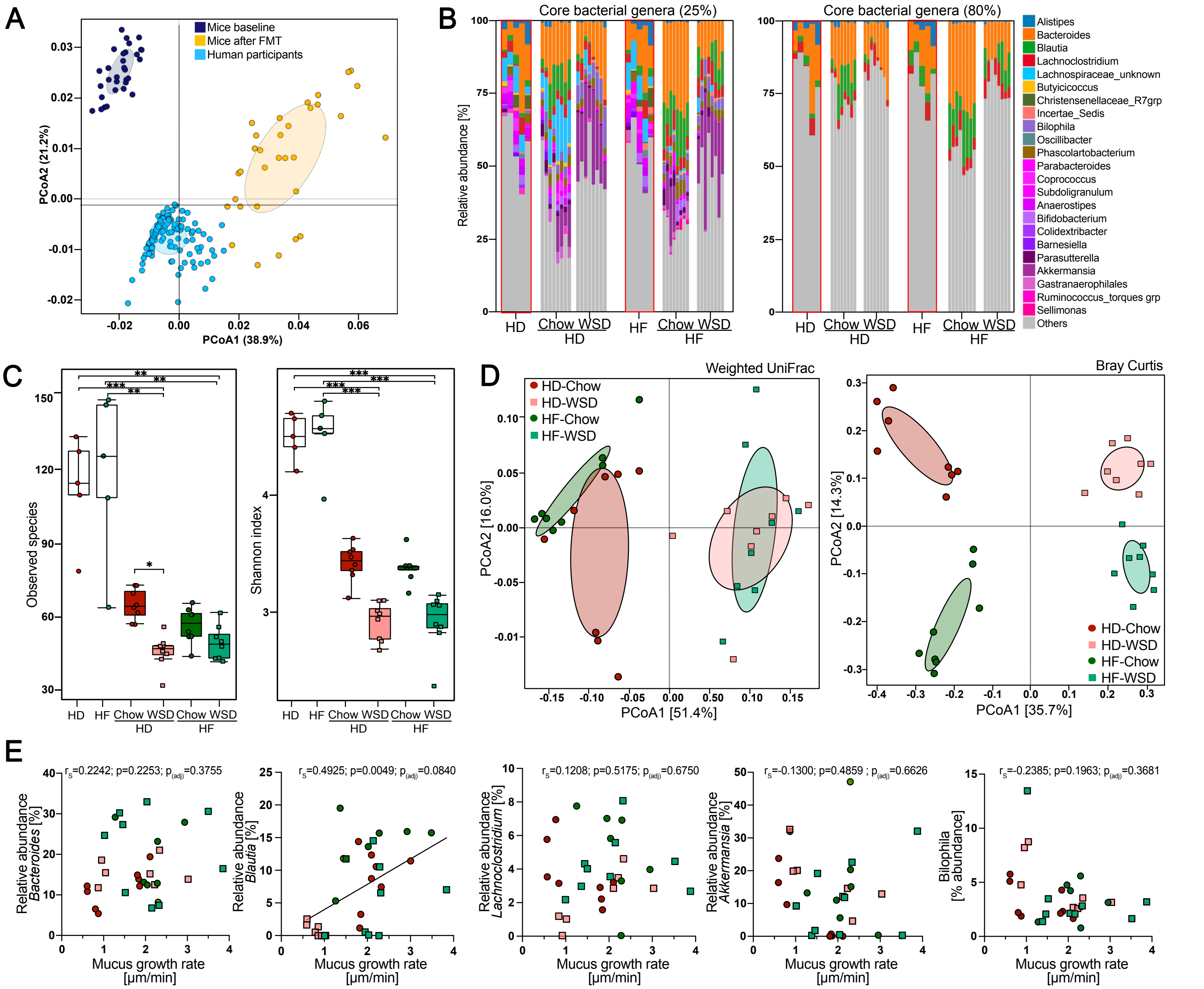

*Supplementary Figure 2*: (A) Weighted UniFrac principal component analysis of fecal bacteria from human study participants (ref. 32) as well as mice before (baseline) and after human-to-mouse fecal microbiota transplantation (FMT). (B) Core bacterial genera relative abundance plots for human donors (highlighted with red margin: HD= habitual diet, HF= high-fiber diet) and human microbiota-transplanted mice fed a chow or Western-style diet (WSD). Core bacterial genera was defined as the bacterial genera present in 25% (left) or 80% (right) of all human study participants and transplanted mouse samples. (C) Stool bacterial alpha-diversity of the human donors and transplanted mice, measured by observed species and Shannon diversity index. Kruskal-Wallis test was used to test for statistical significance. (D) Bacterial beta-diversity, measured by Weighted UniFrac distance matrix and Bray-Curtis dissimilarity matrix, of the transplanted mice. Statistical significance was tested by PERMANOVA with 999 permutations. (E) Spearman correlation analysis between mucus growth rate in the mouse distal colon and relative abundance of selected genera. P<0.05 (*) and p<0.01 (**) are considered statistically significant. Linked to Figure 2.
