## Supplementary material for "The gut commensal *Blautia* maintains colonic mucus function under low fiber consumption through short-chain fatty acid-mediated activation of Ffar2": S3

### Supplementary Figure 3

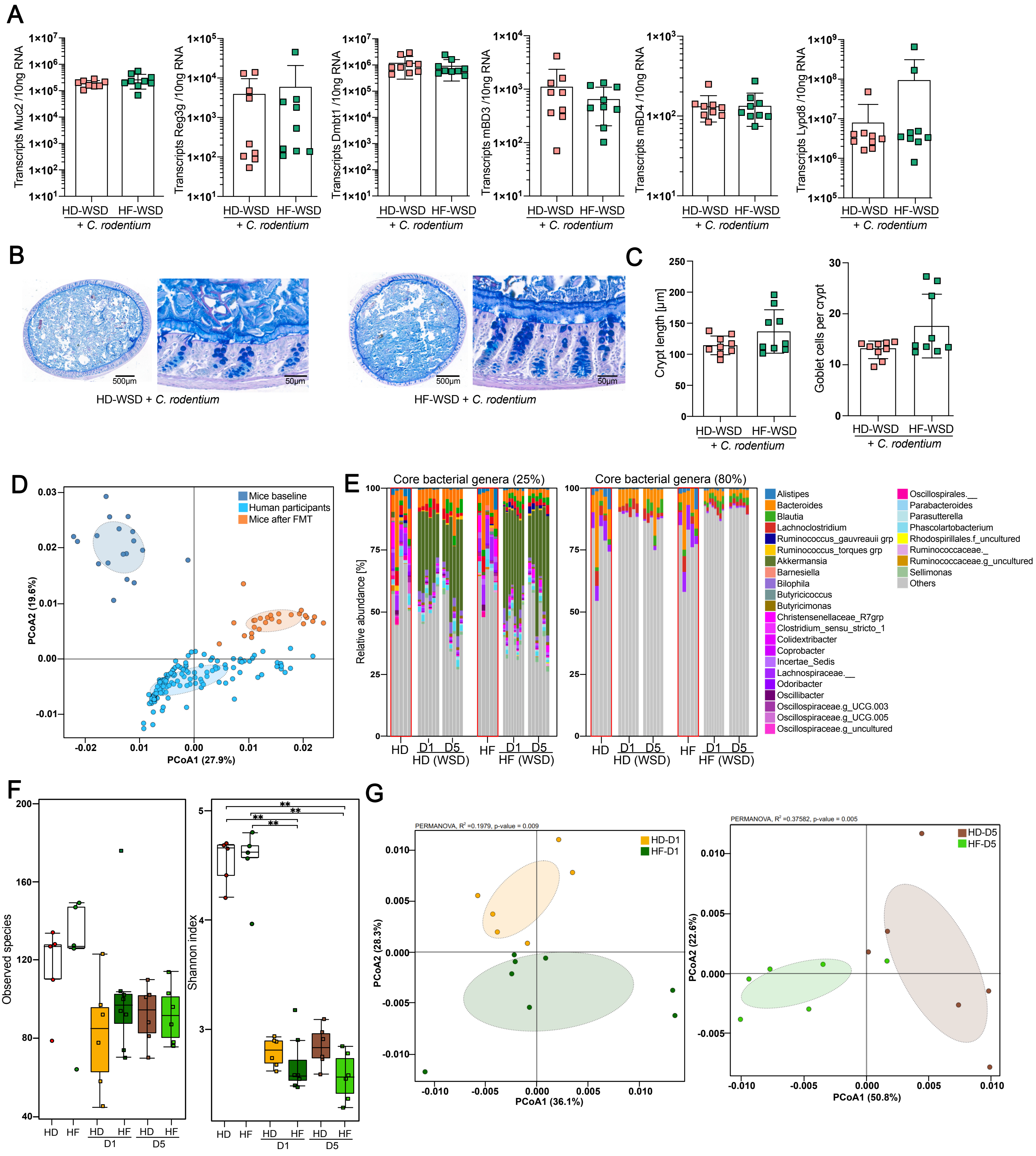


*Supplementary Figure 3*: (A) Absolute quantification of host defense protein/peptide transcripts in the distal colon of mice transplanted with human fecal microbiota and infected with *Citrobacter rodentium*. Human donors consumed their habitual diet (HD) or a high-fiber diet (HF) while mice were fed a Western-style diet (WSD). (B) Alcian Blue/Periodic acid-Schiff (AB/PAS) staining of distal colon sections from the transplanted and infected mice. Representative images from 9 mice/group. Scale bars = 500 µm (full cross-section) and 50 µm (mucosal magnification). (C) Average crypt length and number of goblet cells per crypt in the distal colon of transplanted and infected mice. (D) Weighted UniFrac principal component analysis of bacterial communities from human study participants (ref. 32) and mice before (baseline) and after FMT+*C. rodentium* infection. (E) Core bacterial genera relative abundance plots for human donors (highlighted with red margin: HD= habitual diet; HF= high-fiber diet) and mice transplanted with the human microbiota and infected with *C. rodentium*. (D1=Day 1 post infection; D5=Day 5 post infection). Core bacterial genera was defined as the bacterial genera that were present in 25% (left) or 80% (right) of all human study participants and the transplanted mouse samples. (F) Stool bacterial alpha-diversity of the human donors and the mice transplanted with the human HD and HF samples at D1 and D5 post *C. rodentium* infection, measured by observed species and Shannon diversity index. (G) Weighted UniFrac principal component analysis of stool bacterial genera in the transplanted mice. Normal distribution of the data in A and C was tested with the D'Agostino-Pearson test and statistical significance was tested with an unpaired t-test (normally distributed data) or Mann-Whitney test (non-normally distributed data). Statistical differences in D and G were calculated with PERMANOVA and 999 permutations while Kruskal-Wallis test was used for (F). $p<0.05$ (*), $p<0.01$ (**) and $p<0.001$ (***) are considered statistically significant. Linked to Figure 3.
