## Supplementary material for "The gut commensal *Blautia* maintains colonic mucus function under low fiber consumption through short-chain fatty acid-mediated activation of Ffar2": S4

### Supplementary Figure 4

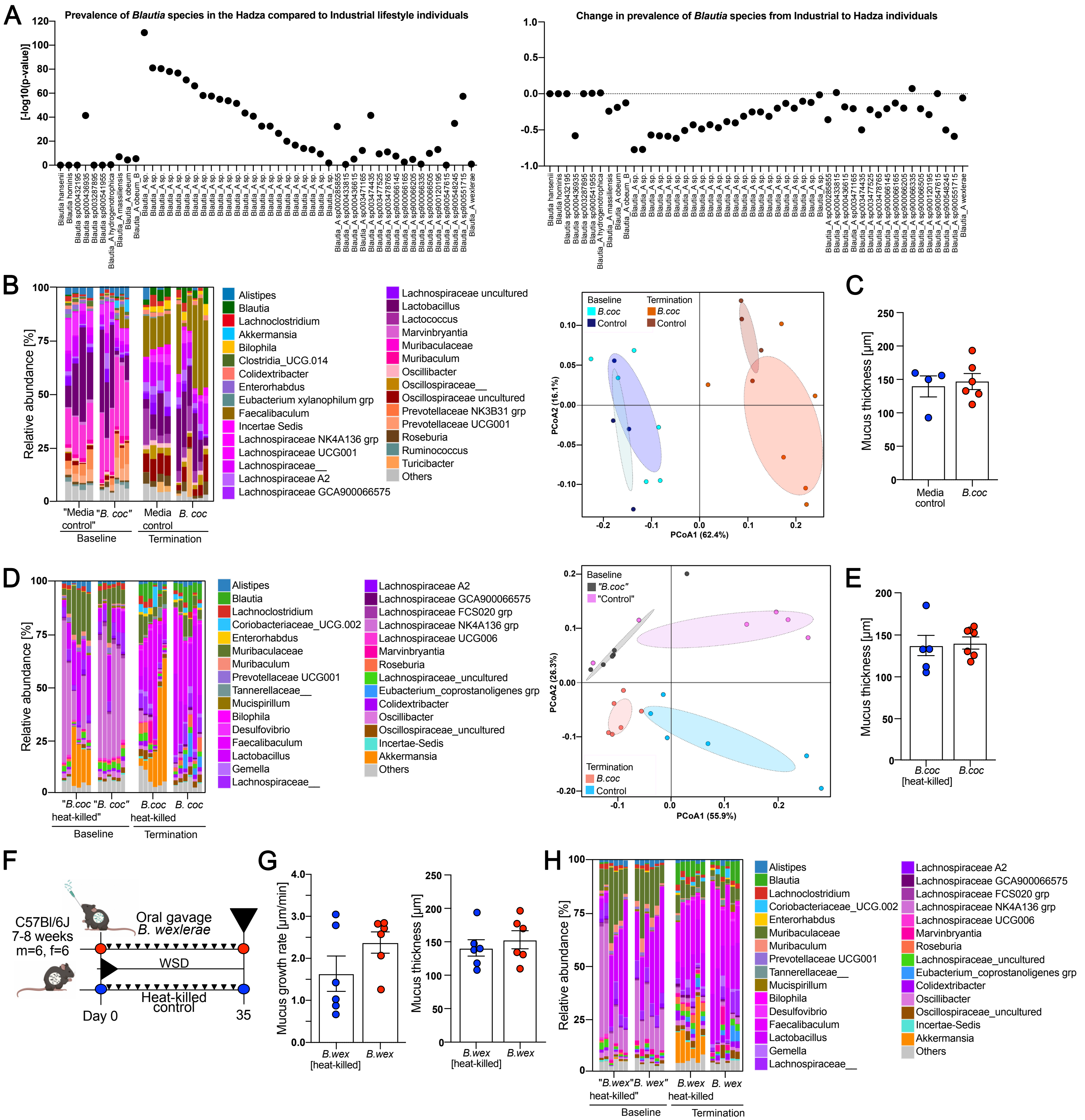


*Supplementary Figure 4*: (A) Comparison of the prevalence of *Blautia* species in the Hadza, modern hunter-gatherers living in Tanzania, compared to industrialized-lifestyle individuals, and changes in *Blautia* prevalence from industrial to Hadza individuals. Left: A log transform of the Fisher's exact test p-value comparing prevalence of the species in the Hadza compared to Industrial lifestyle individuals. Right: The change in prevalence from Industrial to Hadza individuals (value is negative if prevalence is greater in the Hadza). Original data for panel A has been extracted from ref. 37 and is graphically summarized here. (B) Relative abundance of microbial genera and Weighted UniFrac principal component analysis before (baseline) and after (termination) supplementing WSD-fed mice with *B. coccoides* or bacterial growth media through drinking water for 33 days (n=4-6 mice/group). Top 30 bacterial genera are shown. (C) Colonic mucus thickness of the mice supplemented with *B. coccoides* through drinking water. (D) Relative abundance of bacterial genera and Weighted UniFrac principal component analysis before (baseline) and after (termination) supplementing WSD-fed mice with viable or heat-killed *B. coccoides* through oral gavage for 35 days (n=6 mice/group). Top 30 bacterial genera are shown. (E) Colonic mucus thickness of the mice supplemented with live or heat-killed *B. coccoides* through oral gavage. (F) Schematic representation of *B. wexlerae* supplementation through oral gavage to mice fed a WSD. Mice were supplemented with viable or heat-killed *B. wexlerae* (n=6 mice/group) through oral gavage for a period of 35 days, whereupon mucus function was investigated. (G) Mucus growth rate and mucus thickness of the inner colonic mucus layer measured *ex vivo* on viable tissue with a micromanipulator. (H) Relative abundance of bacterial genera before (baseline) and after (termination) supplementing mice fed a WSD with viable or heat-killed *B. wexlerae* through oral gavage for 35 days (n=6 mice/group). Top 30 bacterial genera are shown. Normal distribution of the data in C, E and G was tested with the D'Agostino-Pearson test and statistical significance was tested with an unpaired t-test (normally distributed data) or Mann-Whitney test (non-normally distributed data). Statistical differences in the Weighted UniFrac principal component analyses in B and D were calculated with PERMANOVA and 999 permutations. $p<0.05$ (*), $p<0.01$ (**) and $p<0.001$ (***) are considered statistically significant. Linked to Figure 4.
