## Supplementary material for "The gut commensal *Blautia* maintains colonic mucus function under low fiber consumption through short-chain fatty acid-mediated activation of Ffar2": S5

### Supplementary Figure 5

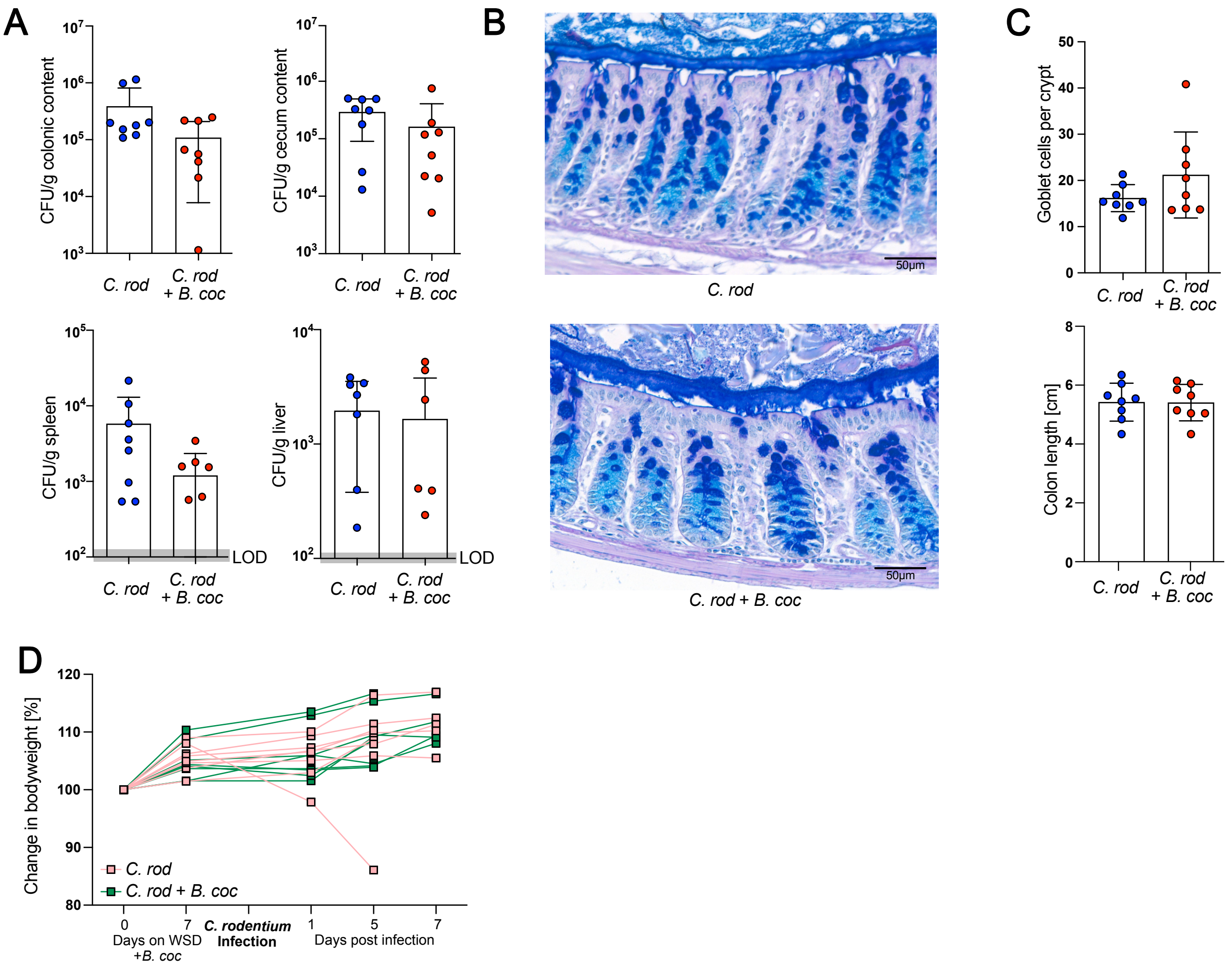


*Supplementary Figure 5*: (A) Colony-forming units (CFU) of *Citrobacter rodentium* in colonic content, cecum content, spleen and liver of mice 7 days after infection, and with or without supplementation of *B. coccoides* (LOD = limit of detection). (B) Alcian Blue/Periodic acid-Schiff (AB/PAS) staining of distal colon sections from mice infected with *C. rodentium*, with or without supplementation of *B. coccoides*. Representative images from 8 mice/group are shown. Scale bars = 50µm. (C) Average number of goblet cells per crypt and colon length in the infected mice. (D) Change in bodyweight before and after *C. rodentium* infection (n = 5-8mice/ group). Data in A and C are presented as means ± SD. Normal distribution in A,C and D was tested with the D'Agostino-Pearson test, statistical significance was determined using an unpaired t-test (normally distributed data) or Mann-Whitney U test (non-normally distributed data) with p< 0.05 (*) considered statistically significant. Linked to Figure 5.
