## Supplementary material for "The gut commensal *Blautia* maintains colonic mucus function under low fiber consumption through short-chain fatty acid-mediated activation of Ffar2": S6

### Supplementary Figure 6

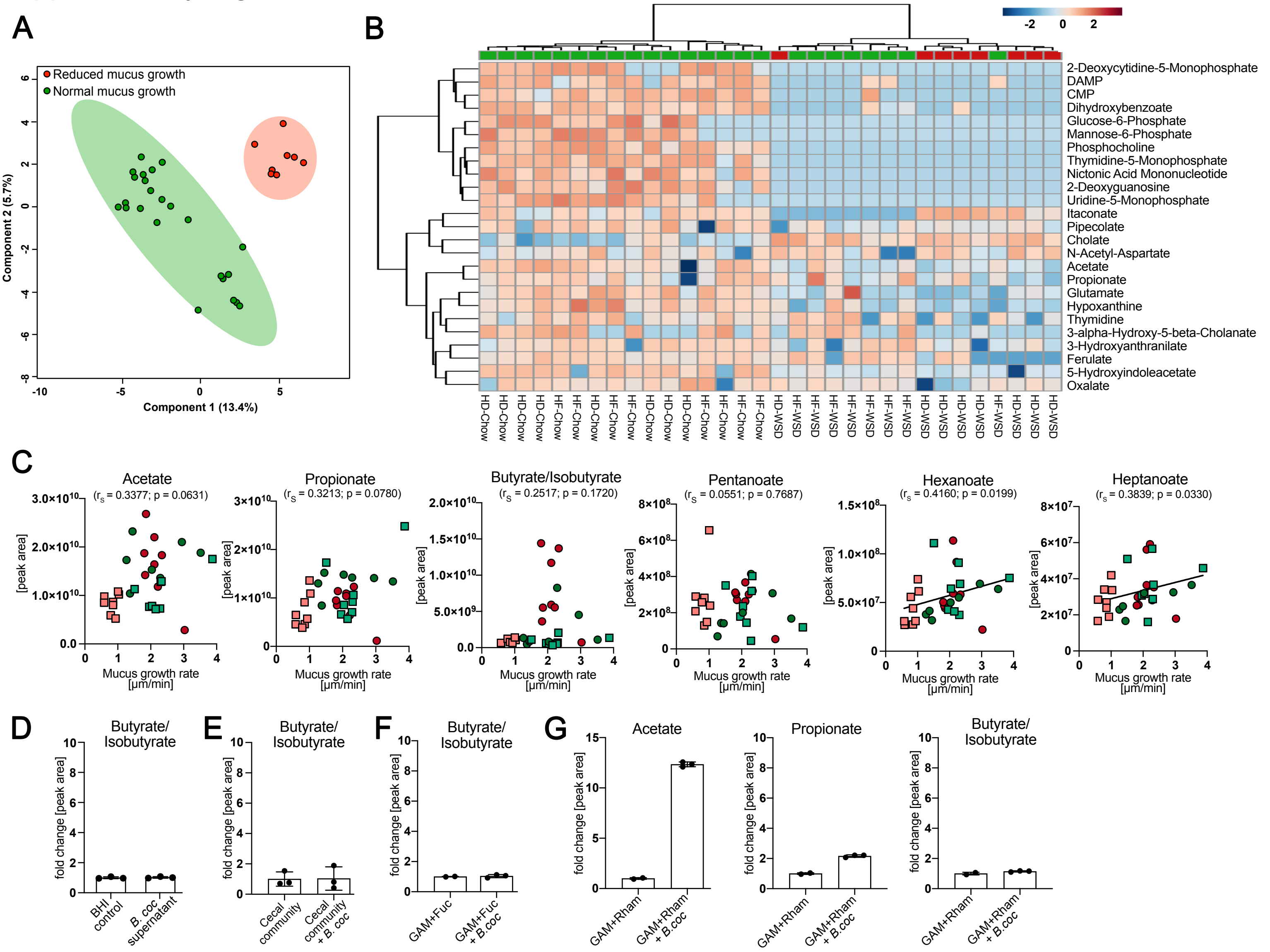


*Supplementary Figure 6*: (A) Partial least squares-discriminant analysis (PLS-DA) of high-throughput global metabolomics profiling of cecal content metabolites. Mouse groups (n=8 mice/group) with average mucus growth rate (HD-Chow, HF-Chow and HF-WSD) are colored in green (n=24) while the mouse group with reduced mucus growth rate (HD-WSD) is colored in red (n=8). (B) Unsupervised hierarchical cluster analysis using Euclidian distance measurement of the 25 most altered metabolites between mucus phenotypes. Color scale indicates fold-change. (C) Correlation analysis between mucus growth rate in the distal colon and peak intensity of the SCFAs, hexanoate and heptanoate, in the cecum of mice transplanted with the human microbiota. Group-specific colour/form coding as in Figure 1. (D) Quantification of butyrate/isobutyrate from the supernatant of a 24h *B. coccoides* culture, the supernatant of a 24h cecal community supplemented with *B. coccoides* (E) and from the supernatant of a 48h B. coccoides culture incubated in GAM media in the presence of fucose (F). (G) Quantification of acetate, propionate and butyrate/isobutyrate from the supernatant of a 48h *B. coccoides* culture incubated in GAM media in the presence of rhamnose. Normal distribution of the data was tested with the D'AgostinoPearson test and data are presented as means ± SD. Statistical significance between the two groups was determined by Mann-Whitney U test (D) while Spearman correlation analysis was used in (C). p<0.05 (*) was considered statistically significant. Linked to Figure 6.
