## Supplementary Table 1 for "The gut commensal *Blautia* maintains colonic mucus function under low fiber consumption through short-chain fatty acid-mediated activation of Ffar2"

| ASV | Potential species |
|---|---|
| 215eab43672099dab90607033e9446f8 | *Blautia coccoides, Blautia pseudococcoides, Blautia hominis, Blautia hansenii, Blautia marasmi* |
| 224b0773460ed546fadcf609add6a4c9 | *Blautia coccoides, Blautia pseudococcoides, Blautia hominis, Blautia hansenii* |
| 64644dbd2c4d66c6dee31dd8a85a98ee | *Blautia coccoides, Blautia pseudococcoides, Blautia hominis, Blautia hansenii, Blautia marasmi* |
| ba11f2b6a318ccd52353476debe4febb | *Blautia coccoides, Blautia hansenii, Blautia producta* |
| 25cf8a8d6aa065f5a5331a1a53db4ba0 | *Blautia coccoides, Blautia hansenii, Blautia producta* |
| 5f371e174ca1193fd938a9ec347c5d48 | *Blautia intestinalis, Blautia sp.* strain Marseille |
| a638c4c026ba6f86f8a31fdefd11d281 | *Blautia wexlerae* |
| 14ec6f561033c1df88c4dfe9de7cb89b | *Blautia wexlerae, Blautia sp.* strain Marseille |
| 4114e476e5cb8dfb4087ff72294f3b49 | *Blautia wexlerae, Blautia sp.* strain Marseille |
| b19e4cfacad1b8143a863bbd394eb705 | *Blautia faecis, Blautia sp.* strain Marseille |
| 91c177085257768127d1752ad41131ad | *Blautia faecis* |
| 891a7b1d20f52c223cb3f898281ba120 | *Blautia sp.* strain Marseille |
| 6a7fd5cbd1a424f24443d9a73c81e579 | *Blautia faecis, Blautia sp.* strain Marseille |
| 16a5f67e3dd348b3f871648a32806e9a | *Blautia caecimuris* |
| 59aa338119796a4be404feda2c098d3b | *Blautia caecimuris* |
| 828bdbef43706f3716448118e0b35dbd | *Blautia obeum* |
| 3c1f95d05561a4329d1e6538d08f424c | *Blautia massiliensis* |

*Supplementary Table 1*: Amplicon sequence variants (ASVs) of *Blautia*-related sequences (left) and the potential species, based on blast search (right).
